# Dopamine-related striatal neurophysiology is associated with specialization of frontostriatal reward circuitry through adolescence

**DOI:** 10.1101/2020.06.24.169847

**Authors:** Ashley C. Parr, Finnegan Calabro, Bart Larsen, Brenden Tervo-Clemmens, Will Foran, Valur Olafsson, Beatriz Luna

## Abstract

Characterizing developmental changes in frontostriatal circuitry is critical to our understanding of adolescent development and can clarify neurobiological mechanisms underlying increased reward sensitivity and sensation seeking, and the emergence of psychopathology during this period. However, the role of striatal neurobiology in the development of frontostriatal circuitry through human adolescence remains largely unknown. We combine longitudinal MR-based assessments of striatal tissue-iron as a correlate of dopamine-related neurobiology with functional magnetic resonance imaging indices of resting-state and reward-state connectivity to investigate the contribution of dopaminergic processes to developmental changes in frontostriatal circuitry. Connectivity between the nucleus accumbens and ventral anterior cingulate, subgenual cingulate, and orbitofrontal cortices decreased through adolescence into adulthood. Nucleus accumbens tissue-iron mediated age-related changes and was associated with variability in connectivity. Our results provide evidence that developmental changes in dopamine-related striatal properties contribute to specialization of frontostriatal circuitry, potentially underlying changes in sensation seeking and reward sensitivity into adulthood.

## Introduction

Adolescence is a period of normative, neurobiological plasticity, marked by heightened sensation seeking and reward-driven behaviors that can be adaptive (Spear, 2000), but can also lead to risk-taking behaviors that undermine survival (e.g., reckless driving, risky sexual behavior, substance use, see Shulman et al., 2016 for review). During this period, adolescents show increased exploratory behavior in novel environments and an increased propensity to seek out experiences motivated by rewarded stimuli (Luciana et al., 2012; Spear, 2000; Stansfield & Kirstein, 2006). These behaviors are thought to be necessary for specializing the neurobiological pathways required to transition into adulthood (Larsen & Luna, 2018; Spear, 2000). Prominent neurodevelopmental models (Shulman et al., 2016) have linked this peak in sensation seeking to maturational changes in frontostriatal reward pathways, which include dopaminergic (DA) projections originating from the ventral tegmental area (VTA) to the nucleus accumbens (NAcc; Kelley & Berridge, 2002; Schultz et al., 2000; Wise, 2004), and to several regions of the ventromedial prefrontal cortex (vmPFC; Haber et al., 1995; Haber & Knutson, 2010). These models propose that adolescent peaks in sensation seeking and reward-driven behavior result from a relative predominance of reward systems over cognitive control systems (Shulman et al., 2016), biasing adolescent decision-making towards rewarding stimuli.

In further support of a DAergic basis for adolescent peaks in sensation-seeking, studies using animal models have shown an increase in striatal DA availability during analogous developmental periods (Andersen et al., 1997; for review, see Luciana et al., 2012). A functional role for NAcc DA in reward seeking behavior during adolescence has been established in animal models (Ernst et al., 2005; Galvan et al., 2006; Luciana et al., 2012; Wahlstrom et al., 2010), and human functional magnetic resonance imaging (fMRI) studies have corroborated a peak in NAcc reward sensitivity in adolescence relative to children and adults (Braams et al., 2015; Douglas et al., 2003; Galvan et al., 2006; Geier et al., 2010; Luna et al., 2013; Padmanabhan et al., 2011; Paulsen et al., 2015, but see Bjork et al., 2004, 2010, who found less NAcc activation in adolescence relative to adults). Adolescent peaks in DA processing are thought to mediate adolescent reward-seeking behaviors through DA’s actions in frontostriatal circuits (Luciana et al., 2012), which are simultaneously undergoing structural and functional specialization through adolescence (Van Duijvenvoorde et al., 2019). Unique developmental patterns of functional connectivity within the mesolimbic pathway have been characterized throughout adolescence, including heightened VTA – NAcc functional connectivity in a rewarded context (Murty et al., 2018), and heightened vmPFC – NAcc resting-state connectivity during adolescence that decreased into adulthood (Fareri et al., 2015; Van Duijvenvoorde et al., 2019; but see Van Den Bos et al., 2012 who saw increased ventral striatal – mPFC connectivity between late childhood through early adulthood when receiving positive incentives). Together, these studies may point to a potential circuit-level mechanism by which reward-seeking behaviors are enhanced in adolescence (Adamantidis et al., 2011; Koob & Volkow, 2010; Luciana et al., 2012), however, the mechanisms linking developmental changes in DA-related striatal function to changes in frontostriatal circuits devoted to reward processing through human adolescence remain unknown.

Elucidating the role of striatal DA-related function in the developmental specialization of frontostriatal pathways during human adolescence has been limited by restrictions in the use of positron emission tomography (PET) in pediatric populations that would allow for *in vivo* indices of DA-function. Addressing these limitations, we used a quantitative MRI-based measure, R2’ (Haacke et al., 2005), to longitudinally assess striatal tissue iron concentration across adolescence as a measure associated with DA neurobiology (Larsen, et al., 2020; Luna et al., 2015; Larsen et al., 2020). Tissue iron has been linked to a number of fundamental neural processes, including cellular respiration (Ward et al., 2014), myelination production and maintenance (Connor & Menzies, 1996; Todorich et al., 2009), and monoamine synthesis (Lu et al., 2017; M. B. H. Youdim & Green, 1978; Youdim, 2018), notably including DA synthesis. Tissue iron is a co-factor for tyrosine hydroxylase (Ortega et al., 2007; Zucca et al., 2017), the rate limiting step in DA synthesis, is located in the dendrites of DA neurons (Torres-Vega et al., 2012), and co-localizes with DA vesicles (Ortega et al., 2007). Moreover, although tissue iron is found throughout the brain, it is highest in the basal ganglia and midbrain (Brass et al., 2006; Hallgren & Sourander, 1958; Thomas et al., 1993). More recently, Larsen et al. (2020) showed cross-sectional and longitudinal correspondence between NAcc tissue iron (R2’) and PET [^11^C]dihydrotetrabenazine (DTBZ), a measure of presynaptic vesicular DA storage (Kilbourn, 2014). Together, these findings indicate that tissue iron reflects important aspects of DA-related striatal neurobiology, and may be an indirect correlate of presynaptic DA function. Tissue iron can be measured non-invasively using MRI, and MR-based assessments of tissue iron have been utilized as an indicator of striatal neurobiology in DA-related disorders, including Parkinson’s disease (Piao et al., 2017; Zucca et al., 2017), Huntington’s disease (Ward et al., 2014), ADHD (Adisetiyo et al., 2014), restless leg syndrome (Allen & Earley, 2007; Connor et al., 2011; Khan et al., 2017), and cocaine addiction (Ersche et al., 2017). Initial studies have characterized normative developmental increases in striatal tissue iron through adolescence that stabilize into adulthood (Larsen, et al., 2020; Larsen & Luna, 2015; Peterson et al., 2019), and more recently, have demonstrated an association between striatal tissue iron concentration and developmental trajectories in cognitive function through adolescence (Larsen, et al., 2020). While the mechanisms underlying this association remain unclear, it is possible that striatal tissue iron affects developmental trajectories in cognitive function, in part, by influencing frontostriatal network organization throughout adolescence. Here, we specifically tested whether normative variation in NAcc tissue iron, given its role in DAergic processes, is related to the development of frontostriatal connectivity through human adolescence.

In order to address this question, we designed a longitudinal developmental neuroimaging study that combines functional magnetic resonance imaging (fMRI) indices of functional connectivity with MR-based assessments of tissue iron (R2’) to probe the role of striatal DA-related neurobiology in developmental changes in frontostriatal connectivity. Given the relevance of this pathway in reward processing and decision-making (Cools, 2011; Knutson, Adams, et al., 2001; Knutson et al., 2001; Schultz, 2002; Wise, 2004), we examined functional connectivity during a rewarded-state in which participants learned the value of actions on the basis of rewards and translated these value representations to instrumental actions, in addition to resting-state connectivity. We then investigated the extent to which *in vivo* indices of NAcc neurobiology (R2’) mediated developmental changes in vmPFC – NAcc connectivity through adolescence.

We show developmental decreases in vmPFC – NAcc connectivity through adolescence, confirmed in both reward-state and resting-state contexts. Further, we find that NAcc R2’ is associated with individual differences in the strength of vmPFC – NAcc connectivity, and significantly mediates observed age-related decreases in connectivity. Together, these results provide strong evidence that developmental changes in DA-related striatal neurobiology may play a key role in the specialization of frontostriatal circuitry during adolescence, providing a potential mechanism for decreased sensation seeking into adulthood.

## Materials and Methods

### Participants

One hundred and forty-four adolescents and young adults participated in an accelerated longitudinal study that included behavioral and MRI sessions (76 females, mean age, 19.5, age range, 12 – 31). Ninety-nine participants also participated in a follow-up visit approximately 18 months following the first session (47 females, mean age, 22.0, age range, 13 – 33). A total of 243 sessions were included. Participants were recruited from the community and were screened for the absence of psychiatric and neurological problems including loss of consciousness, self or first-degree relatives with major psychiatric illness, and contraindications to MRI (e.g., claustrophobia, metal in the body, pregnancy). Participants or the parents of minors gave informed consent with those less than 18 years of age providing assent. All experimental procedures were approved by the University of Pittsburgh Institutional Review Board and complied with the Code of Ethics of the World Medical Association (Declaration of Helsinki, 1964). Participants 18 years of age and older underwent simultaneous PET acquisition, and previous publications have been reported on this line of inquiry (Calabro et al., *in preparation*; Larsen et al., 2020). Here, given that the current research question was developmental in nature, we included MR-based indices that were collected across the entire adolescent sample.

### Reward-Guided Decision-Making Task

To characterize brain connectivity in a rewarded context, we used fMRI data acquired during a reward learning task (Fig. 1A). Briefly, participants completed six 5-minute blocks during which they were required to choose a cell from a 3×3 grid map where each map location was pre-determined to have a randomized reward probability of 20%, 50%, or 80% (Fig. 1B). On each trial, participants were presented with the option to move to one of two different map locations. The two locations were initially represented by hash marks (“#”) for a pseudorandom duration of 1-6 seconds (“expectation and decision-making” epoch), after which the symbols changed to display “1” and “2”. Participants then made their response, and the avatar moved to reflect their new location on the map. Visual and auditory feedback was then presented to indicate whether they received a reward or not for that trial (“feedback” epoch), based on the chosen location’s predetermined reward probability, and if so, whether it was a low reward (single coin, 75% of rewarded trials) or a high reward (multiple coins, 25% of rewarded trials). Accuracy on this task was calculated as the proportion of trials in which participants correctly selected the map location with the higher reward probability.

**Figure 1.**
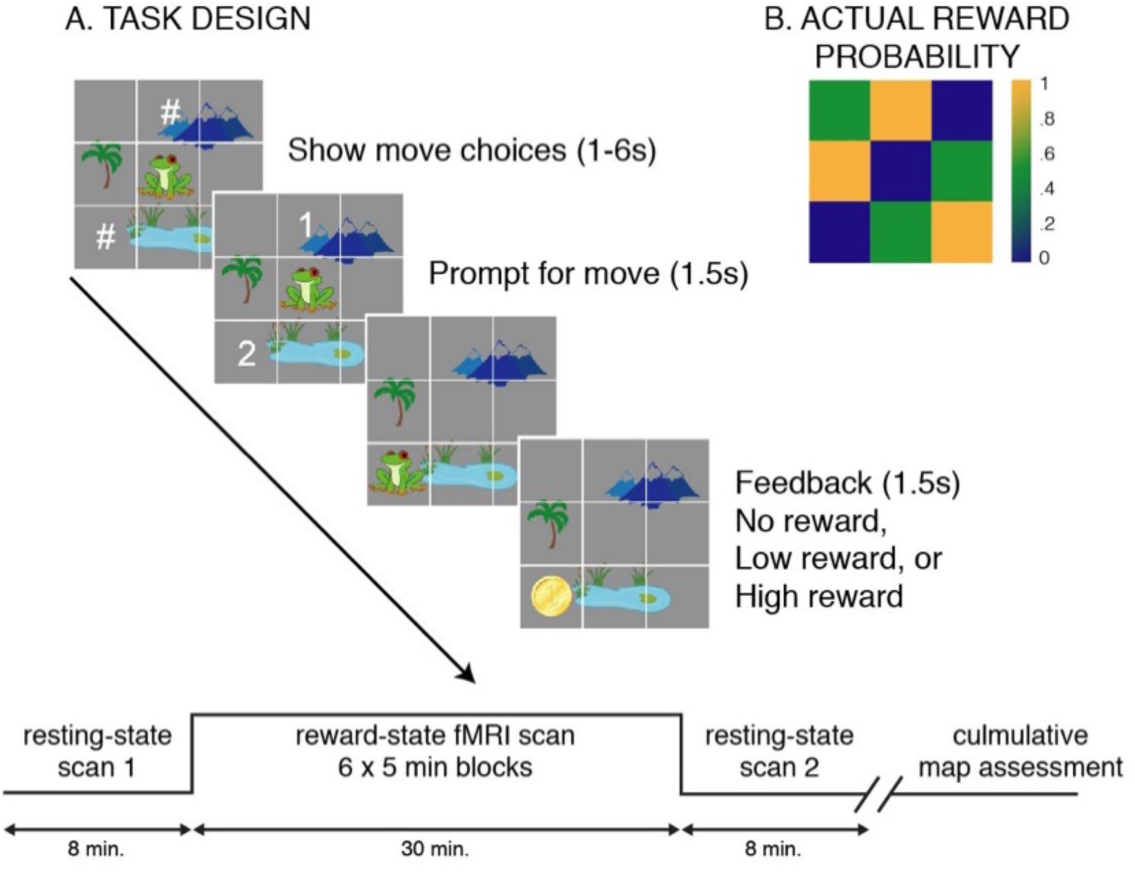
Reward task design. (A). Reward map learning task. Participants were presented the option to choose one of two different map locations and were rewarded based on the chosen location’s predetermined reward probability and magnitude. (B) Actual reward probabilities. Map represents actual reward probabilities associated with each square in (A). High probability squares are represented in yellow, medium in green, and low in blue.

The underlying probabilities were maintained throughout the six task runs, and participants were instructed to learn probability contingencies in order to maximize their earnings (up to $25) over the course of the session. The actual reward sequence was matched across participants to ensure consistency in the pattern of positive and negative feedback, such that all participants earned the maximum reward, though they were unaware of this at the time. Participants were presented overall feedback on their progress (total earnings, in arbitrary unit points) at the end of each block, and feedback about total monetary earnings was given after the entire scan was completed.

Finally, to assess learning, participants performed a ‘Map Test’ after the scan where they were sequentially presented with 50 pairs of map locations and were instructed to recall and report which of the two squares were associated with higher rewards when they had performed the task in the scanner.

In order to learn the mechanics of the task, participants were trained using an abbreviated (single 5-minute block) version of the task during both the behavioral session occurring prior to the scan day, and immediately prior to entering the scanner. Participants were informed that the maps used during the training sessions would be different from the map used during the scan.

On a portion of trials, after receiving feedback, participants performed “bonus trials” consisting of either a rewarded or unrewarded anti-saccade task (described previously in Geier et al., 2010; Murty et al., 2018; Ordaz et al., 2013). Although these data were modeled in the fMRI analyses described below, data is not presented here.

### MR Data Acquisition & Preprocessing

MRI and adult PET data (18 – 31-year-old cohort only) were collected simultaneously over 90 minutes on a 3T Siemens Biograph mMR PET/MRI scanner. Participants’ heads were immobilized using pillows placed inside the head coil, and participants were fitted with earbuds for auditory feedback and to minimize scanner noise. Structural images were acquired using a T1 weighted magnetization-prepared rapid gradient-echo (MPRAGE) sequence (TR, 2300 ms; echo time (TE), 2.98 ms; flip angle, 9°; inversion time (T1), 900 ms, voxel size, 1.0 × 1.0 × 1.0 mm). Functional images were acquired using blood oxygen level dependent (BOLD) signal from an echoplanar sequence (TR, 1500 ms; TE, 30 ms; flip angle, 50°; voxel size, 2.3 × 2.3 mm in-plane resolution) with contiguous 2.3mm – thick slices aligned to maximally cover the cortex and basal ganglia. Thirty minutes (six 5-min blocks) of reward-context functional connectivity was compared to 16 minutes (two 8 min sessions) of fixation resting state fMRI data collected prior to (pre-task) and following (post-task) performing the task.

Structural MRI data were preprocessed to extract the brain from the skull, and warped to the MNI standard brain using both linear (FLIRT) and non-linear (FNIRT) transformations. Resting-state data were preprocessed using a pipeline that minimized the effects of head motion (Hallquist et al., 2013) including 4D slice-timing and head motion correction, skull stripping, intensity thresholding, wavelet despiking (Patel et al., 2014), coregistration to the structural image and nonlinear warping to MNI space, local spatial smoothing with a 5mm Gaussian kernel based on the SUSAN algorithm, intensity normalization, and nuisance regression based on head motion (6° of translation/rotation and their first derivative) and non-gray matter signal (white matter and CSF and their first derivative). Bandpass filtering between .009 and .08 Hz was done simultaneously with nuisance regression.

To measure functional connectivity within a rewarded context, we applied a background connectivity approach (Al-Aidroos et al., 2012; Fair et al., 2007; Norman-Haignere et al., 2012; Salvador et al., 2005) where task-related trial level BOLD responses of the time series are regressed out using a general linear model (GLM; (Dosenbach et al., 2010; Elliott et al., 2019; Fair et al., 2007; Greene et al., 2018) revealing the BOLD fluctuations that are sustained throughout a task. The six 5 min runs were concatenated, providing 30 minutes of data to optimally estimate and remove task-evoked BOLD responses. Task-evoked events were modeled with a TENT function to estimate the hemodynamic response for 24 s after each event in 1.5 s increments (using 3dDeconvolve in AFNI; Murty et al., 2018). In comparison to the canonical hemodynamic response function (HRF), the TENT function makes no assumptions about the shape of HRF responses, and should therefore remove more variability due to task-evoked signals from the time series (Chen et al., 2015).

We modeled the following task events: the hash mark interval (“expectation and decision-making” epoch, Fig. 1A) and the feedback event (in which participants made their selection and received feedback, Fig. 1A). Feedback events were modeled separately for no reward, low reward, and high reward trials. We additionally modeled the preparatory and outcome phases of the anti-saccade task using a canonical (“GAM”) hemodynamic response function, based on our previous investigations of this task (Geier et al., 2010). The residuals from this model (i.e., the error term from the GLM) were then preprocessed in a similar manner as the resting state data, including bandpass filtering between .009 and .08 HZ using AFNI’s 3dTproject with NTRP censor to match the temporal characteristics of the resting-state data.

Frame-wise motion estimates were computed for both resting state and background data, and volumes containing frame-wise displacement (FD) >0.3mm were excluded from analyses (Siegel et al., 2014). Participants with more than 25% of TRs censored were excluded from analysis. Importantly, the total number of censored volumes was not associated with participant age (*β*= .002, *t* = .92, *p* = .36).

### Acquisition and preprocessing of R2’ data

R2’ reflects the reversible transverse relaxation rate (1/T2’), which is the difference between the effective (R2*; 1/T2*) and irreversible (R2; 1/T2) relaxation rates (Haacke et al., 2005). R2 data were acquired using multi-echo turbo spin echo (mTSE; effective echo times 12, 86, and 160ms; TR = 6580ms; 12ms spacing between spin refocusing pulses; FoV = 240×240 mm^2^; 27 3mm transverse slices; 1mm slice gap). R2* data were acquired using multi-echo gradient echo (mGRE; echo times 3, 8, 18, and 23ms; TR = 724ms; flip angle, 25°; FoV = 240×240 mm^2^). R2 and R2* images were registered to MNI space, and estimates of R2’ were derived for each participant by subtracting R2 from R2* estimates (detailed methodologies for calculating R2’ are described in Larsen et al., 2020). All R2’ images were visually assessed by authors B.L. and V.O., and confirmed by A.P. and other independent raters for the presence of motion, shimming, or registration artifacts. Data quality was rated on a five-point scale for artifact intrusion on striatal signal, in addition to visible macroscopic field inhomogeneities affecting the striatum. Participants with scores of three out of five or above were excluded, resulting in 66 sessions not being included in R2’ analyses.

In order to control for age related accumulation of tissue iron (Aquino et al., 2009; Hallgren & Sourander, 1958; Wang et al., 2012), we regressed age from the R2’ measure with an inverse age function (age^-1^), with R2’ as the dependent variable, age^-1^ as the fixed-effects factor. The residuals of this model (age-residualized R2’) were included in the analyses detailed below as an index of striatal dopamine neurobiology (i.e., relative R2’ level for a participant’s age). Extreme value outliers were detected as residuals that exceeded three standard deviations from the mean after removing age-effects and were not included in the final regression models. Exclusion based on R2’ quality assessment and outlier detection was not significantly associated with participant age (β = -0.02, t = 0.86, p = .39). Following quality assessments, R2’ indices of striatal physiology were obtained in 105 12-31year old participants (53 female) at visit 1, of which 72 returned approximately 18 months later for a second session (31 female), for a total of 177 sessions. R2’ data from a subset of this cohort has been presented in a previous publication (Larsen et al., 2020).

### Statistical analysis of behavioral data

#### Age

To evaluate the relationship between age and map test performance, we implemented linear mixed-effects models (RStudio version 1.1.463; Bates et al., 2014), including random intercepts estimated for each participant in order to account for the longitudinal aspect of the dataset. We tested linear and inverse (age^-1^) functional forms of age and selected the model with the lowest Akaike Information Criterion (AIC), indicating best model fit. In the case that the AIC for each model was equivalent, we characterized developmental changes with an inverse age function, as we hypothesize that development is characterized by an initial period of rapid growth followed by stabilization into adulthood, and previous studies have shown that inverse age provides a better fit for developmental changes through adolescence (Luna et al., 2004; Murty et al., 2018; Ordaz et al., 2013; Simmonds et al., 2017). Sex and visit number were included as covariates, to ensure the robustness of the associations. We tested for sex by age interactions, and the interaction term was removed from the final model as it was not significant.

#### R2’

We investigated the effects of age-residualized R2’ on performance using linear mixed-effects models, including random intercepts estimated for each participant. This analysis was repeated while including sex and visit number as covariates. Again, we first tested for sex by age interactions, and this interaction term was removed from the final model as it was not significant.

### Statistical analysis of fMRI data

#### Functional Connectivity Analyses

To investigate functional connectivity between the vmPFC and the NAcc, we implemented two separate analysis strategies. First, we used a region of interest (ROI) approach investigating connectivity between hypothesis-driven, predefined a priori ROIs (described in detail below). Next, to ensure our ROI selection did not bias our results, we conducted exploratory whole-brain voxelwise connectivity analyses with the nucleus accumbens (NAcc) as the seed region (described in detail in Supplemental Methods).

#### A Priori Region of Interest Selection

Bilateral vmPFC and NAcc ROIs were selected on the basis of their established role within reward and valuation circuitry (Cools, 2011; Knutson, Adams, et al., 2001; Knutson, Fong, et al., 2001; Schultz, 2002; Wise, 2004). Left and right NAcc were defined using the Harvard-Oxford subcortical atlas (Fig. 2A; Jenkinson et al., 2011). The vmPFC was sub-divided into eight bilateral subregions using the Mackey & Petrides vmPFC atlas (Fig. 2B; Mackey & Petrides, 2014). All ROIs included only voxels with a 50% or greater probability of being grey matter in the MNI-152-09c template. ROIs were excluded from analysis if they did not have at least 70% coverage in at least 70% of participants. These criteria resulted in exclusion of the bilateral anterior medial OFC ROIs (areas 14r’ and 11m in Mackey & Petrides, 2014 and dark grey areas in Fig. 2B), as well as the more rostral posterior medial OFC ROI (area 14r in Mackey & Petrides, 2014 and dark grey areas in Fig. 2B). Our final list of ROIs included the following ROIs for the left and right hemisphere: posterior medial OFC (area 14c), anterior vmPFC (area 14m), ventral ACC (area 24), subgenual cingulate (area 25), rostral ACC (area 32), and the NAcc (Fig. 2).

**Figure 2.**
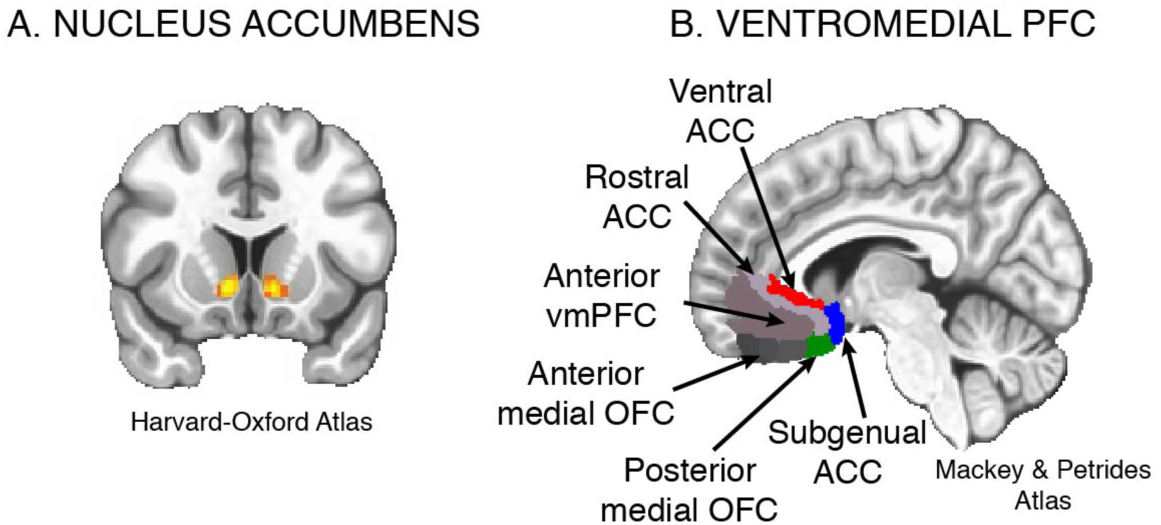
Region of interest selection. (A) Nucleus accumbens was defined using the Harvard – Oxford subcortical atlas (Jenkinson et al., 2011), and (B) Ventromedial prefrontal cortex subregions were defined using the Mackey & Petrides atlas (Mackey & Petrides, 2014). ACC, anterior cingulate cortex, vmPFC, ventromedial prefrontal cortex, OFC, orbitofrontal cortex.

#### ROI Functional Connectivity Analyses

For each ROI, time series for both the reward- and rest-contexts were extracted from each participant by taking the first principal component across all voxels within each ROI. Pearson correlation coefficients were then computed between the NAcc seeds (left and right hemisphere) and the five vmPFC seeds (left and right hemisphere) and normalized using Fisher’s Z transformation.

#### Associations between age and connectivity

Our primary analyses utilized left and right ROI pairs in a single model with hemisphere (i.e., left/right hemisphere of the NAcc) and laterality (i.e., ipsilateral: L->L, R->R; contralateral: L->R, R->L) as within-participants factors (Tervo-Clemmens et al., 2017). Models were run separately for each ROI pair. To assess differences in connectivity as a function of age and context in each ROI pair (main effects), we used linear mixed-effects models with the correlation value as the dependent variable, and context (rest, reward), age, sex, head motion, seed ROI hemisphere, laterality, and visit number as fixed-effects factors; random intercepts were estimated for each participant. As in the behavioral analysis, we tested and selected the best fitting age-model across all ROIs. We next examined differential age effects across contexts (age by context interactions). This interaction term was removed from the final models as it was not significant. Data were considered significant at *p* < .01 to account for multiple comparisons (Bonferroni correction: 0.05/5 connectivity pairs, corrected alpha=.01). The same models were implemented in the whole-brain voxelwise analyses (linear mixed-effects models implemented in AFNI; see Supplemental Methods).

#### Associations between tissue-iron and connectivity

In the connectivity pairs where ROI analyses revealed significant developmental effects, we further interrogated their association with indices of striatal tissue iron (age-residualized NAcc R2’) using linear mixed-effects models while modelling age, sex, context, head motion, seed ROI hemisphere, laterality, and visit number as covariates, and participant ID as the random-effects factor. We next examined differential effects of R2’ on connectivity within each context (R2’ by context interactions), and this interaction term was removed from the final models when not significant, unless otherwise stated. The same models were implemented in the whole-brain voxelwise analyses (linear mixed-effects models implemented in AFNI; see Supplemental Methods).

To test the hypothesis that differences in R2’ indices contributed to age-related changes in connectivity, mediation analyses were performed on each ROI that had significant associations with age and R2’. The mixed-effects models employed in the mediation analysis were implemented as in the previous analyses. Significance values for indirect effects were obtained using 10000 draws in a bootstrap procedure (mediation package in R; Tingley et al., 2014). Participant ID was entered into these models as a random-effects variable to account for the repeated (longitudinal) nature of the dataset.

#### Associations between performance and connectivity

Finally, we examined the association between map test performance and connectivity in a separate model, using the same covariates detailed above. Again, we examined differential effects of performance on connectivity within each context (performance by context interactions), and this interaction term was removed from the final models as it was not significant.

## Results

### Behavioral data

#### Task Performance Improves with Increasing Age

Map test performance improved with increasing age (Fig. 3A; 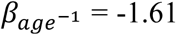, *t* = -2.26, *p* = .02), with inverse age resulting in marginally better model fits of the data (map test performance: linear AIC, -298.1; inverse AIC, -298.9). There was no significant age by sex interaction (*β* = -.004, *t* = -1.24, *p* = .22).

**Figure 3.**
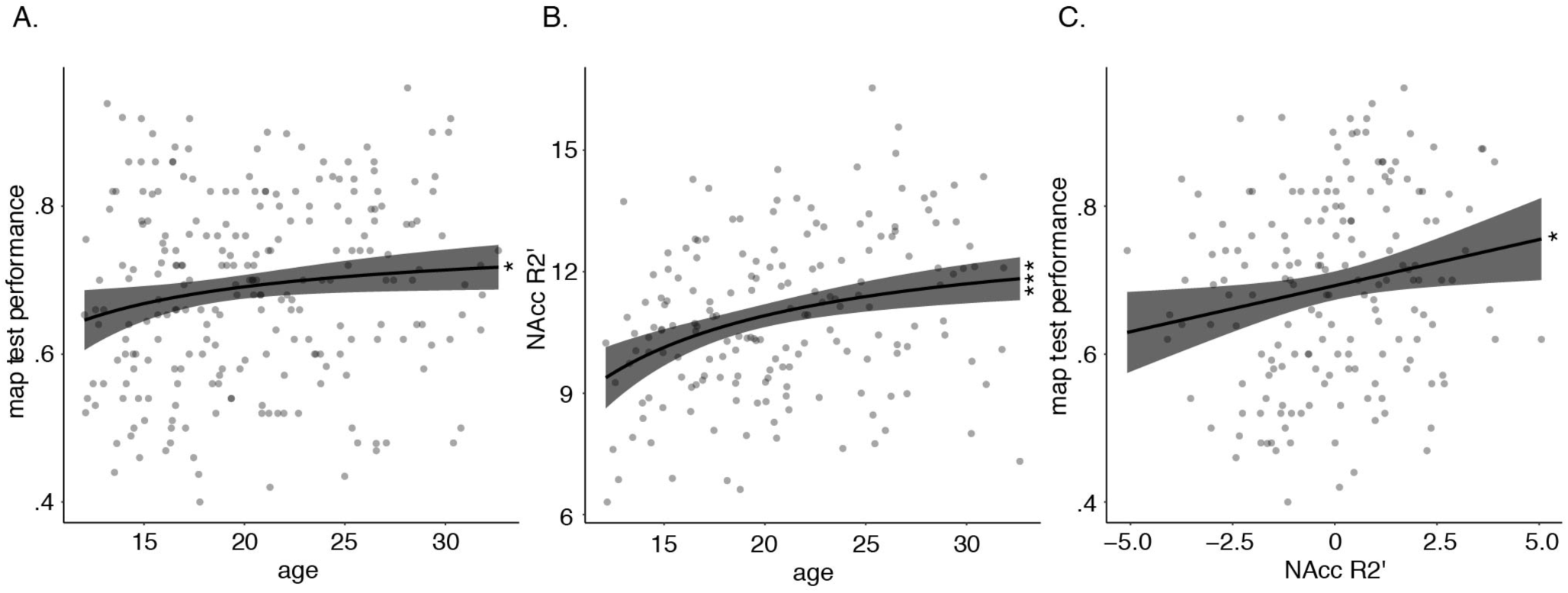
Associations between age, task performance, and nucleus accumbens R2’. (A) Map test performance improves across adolescence. (B) NAcc R2’ increases across adolescence. (C) Higher R2’ (age-residualized) is associated with better map test performance. In (C), the y-axis represents age-residualized estimates of R2’. In (A – C), shaded region represents 95% confidence interval. * *p* _Bonferroni_ < .05, ** *p* _Bonferroni_ < .01, *** *p* _Bonferroni_ < .001.

### fMRI data

#### vmPFC – NAcc Connectivity Decreases Throughout Adolescence

We ran mixed-effects models to investigate connectivity as a function of age- and context-in each ROI pair. We observed a main effect of age on the connectivity of the NAcc with three cortical regions (while covarying for sex, visit number, and motion estimates, see methods): 1. ventral ACC (Fig. 4A; 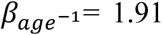, *t* = 3.15, *p* = .002, *p*_Bonferroni_ = .01); 2. subgenual cingulate (Fig. 4A; 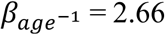, *t* = 3.71, *p* < .001, *p*_Bonferroni_ < .001); and 3. posterior medial OFC (Fig. 4A; 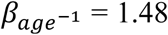, *t* = 2.63, *p* = .008, *p*_Bonferroni_ = .04). In all cases, inverse age resulted in better model fits to the data, and increasing age was associated with significant decreases in connectivity across all 3 ROI pairs in both reward and rest contexts (Fig. 4A). Additionally, these models revealed a significant main effect of context in each of the ROI pairs. While connectivity between the NAcc and ventral ACC was greater during the rest versus reward context (β = 0.11, *t* = 15.69, *p* < .001, *p*_Bonferroni_ < .001), connectivity between NAcc and subgenual ACC pair (β = -0.05, *t* = -6.96, *p* < .001, *p*_Bonferroni_ < .001), and NAcc and posterior medial OFC, was greater during reward context than rest (β = -0.05, *t* = -6.34, *p* < .001, *p*_Bonferroni_ < .001). Notably however, there were no significant context by age interactions for any ROI (all *p*_Bonferroni_ > .05), indicating that the observed age effects were consistent across the two contexts. Therefore, for visualization of overall developmental effects, we have combined connectivity values from both rest- and reward-states in Fig. 4A.

**Figure 4.**
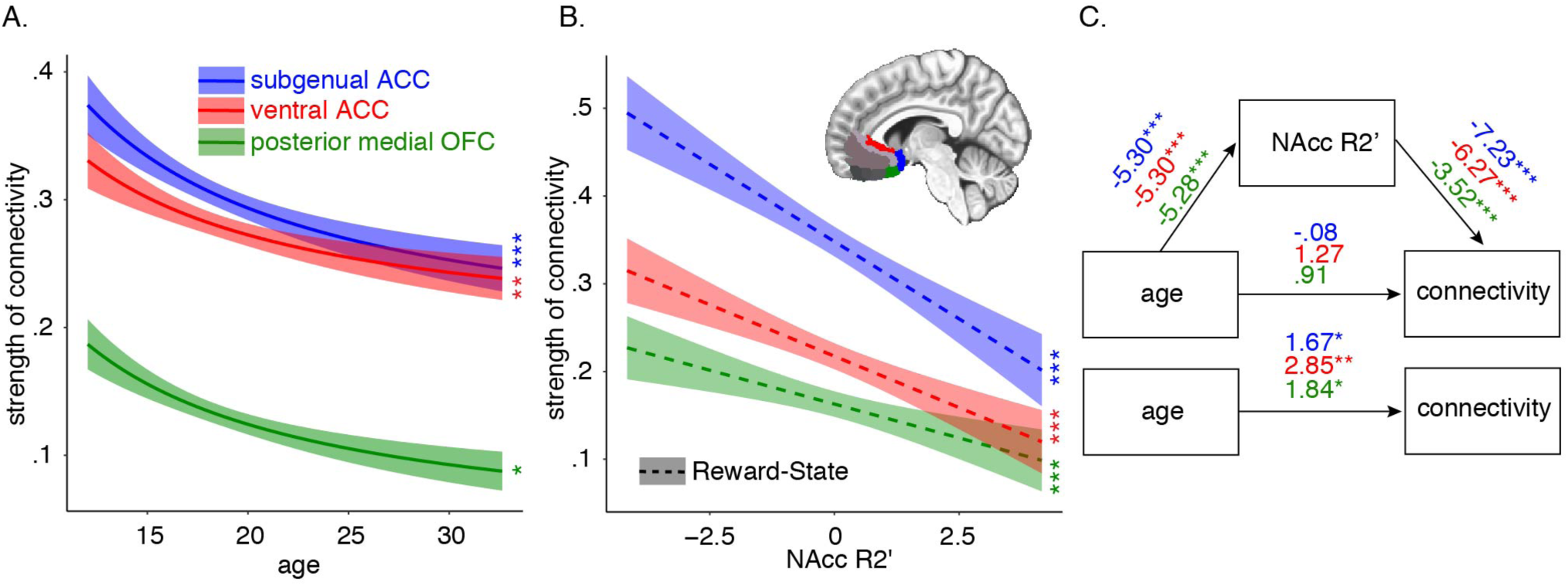
Age-related changes in connectivity and their association with nucleus accumbens R2’. (A) Nucleus accumbens to ventral anterior cingulate, subgenual cingulate, and posterior medial orbitofrontal connectivity decreases through adolescence. (B) Higher nucleus accumbens R2’(age-residuals) is associated with decreased connectivity across all 3 ROI pairs. (C) R2’ significantly mediates the relationship between age and functional connectivity. In (A), resting-state and reward-state are combined. In (B) and (C), we show reward-state connectivity for visualization purposes (see Fig. S1 for resting-state results). In (B), the x-axis represents age-residualized estimates of R2’. In (A – B), shaded region represents 95% confidence interval. * *p* _Bonferroni_ < .05, ** *p* _Bonferroni_ < .01, *** *p* _Bonferroni_ < .001. ACC, anterior cingulate cortex, OFC, orbitofrontal cortex.

#### vmPFC – NAcc Connectivity and Task Performance

Mixed-effects models investigating connectivity as a function of performance during the reward learning task (see methods & supplemental methods) did not reveal any significant main effects of performance (all *p*_Bonferroni_ > .05), any age by performance interactions (*p*_Bonferroni_ > .05), or any context by performance interactions (*p*_Bonferroni_ > .05) in any ROI pair (data not shown).

### Tissue Iron data

#### R2’ Increases with Age

As has been previously shown (Larsen et al., 2020), NAcc R2’ increased with age (Fig. 3B; 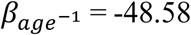, *t* = -3.85, *p* < .001), with inverse age resulting in better model fits of the data (linear AIC, 695.4; inverse AIC, 691.5). There was no significant age by sex interaction (*β* = 2.72, *t* = .11, *p* = .91).

To control for age related accumulation of tissue iron (Aquino et al., 2009; Hallgren & Sourander, 1958; Wang et al., 2012), age-residualized estimates of R2’ were utilized in the subsequent analyses (see methods), and reflect the relative R2’ level for a participant’s age cohort.

#### Task Performance is better in Individuals with Relatively High NAcc R2’

Map test performance was positively associated with NAcc R2’ (Fig. 3C; age residuals; *β* = 0.01, *t* = 2.21, *p* = .03), indicating that individuals with relatively high R2’ *for their age cohort* had better performance relative to those with low R2’ indices. There was no significant R2’ by sex interaction (*β* = -.004, *t* = -.37, *p* = .71).

#### Decreased vmPFC – NAcc Connectivity is Associated with Higher NAcc R2’

Mixed-effect models to investigate connectivity in each ROI pair as a function of R2’ (age-residualized estimates) and context (while covarying for sex, visit number, and motion estimates, see methods) revealed a main effect of R2’ in all three ROI pairs that showed age effects: 1. ventral ACC – NAcc (Fig. 4B & Fig. S1A; *β*= -0.02, *t* = -4.99, *p* < .001, *p*_Bonferroni_ < .001); 2. subgenual cingulate – NAcc (Fig. 4B & Fig. S1A; *β*= -0.02, *t* = -6.17, *p* < .001, *p*_Bonferroni_ < .001); and 3. posterior medial OFC – NAcc (Fig. 4B; *β*= -0.01, *t* = -2.41, *p* = .01, *p*_Bonferroni_ = .05). In all cases, higher indices of R2’ with respect to age were associated with significant decreases in connectivity across all 3 ROI pairs (Fig. 4B & Fig. S1A). Additionally, we observed a significant R2’ by context interaction on subgenual cingulate – NAcc (*β*= 0.02, *t* = 4.73, *p* < .001, *p*_Bonferroni_ < .001) and posterior medial OFC – NAcc connectivity (*β*= 0.01, *t* = 3.07, *p* = .002, *p*_Bonferroni_ = .01). To further characterize the nature of these interactions, we conducted follow up tests within each context. For the sACC – NAcc pair, these tests revealed that R2’ was associated with greater decreases in the reward context (Fig. 4B; *β*= -0.04, *t* = -7.19, *p* < .001, *p*_Bonferroni_ < .001), however, the effect of R2’ on connectivity was still significant during rest (Fig. S1A; *β*= - 0.02, *t* = -3.71, *p* < .001, *p*_Bonferroni_ < .001). A similar pattern was observed for the OFC – NAcc pair, such that R2’ was significantly associated with decreased connectivity in the reward context (Fig. 4B; *β*= -0.02, *t* = - 3.46, *p* < .001, *p*_Bonferroni_ < .001), however here, the relationship between connectivity and R2’ was not significant in the resting-state context (Fig. S1A; *β*= -0.003, *t* = -0.70, *p* = .48, *p*_Bonferroni_ = 1.00). For simplification, we show connectivity values from the reward-state connectivity in Fig. 3B, and resting-state values are shown in Fig. S1A.

#### NAcc Tissue-Iron Mediates Decreases in vmPFC – NAcc Connectivity Throughout Adolescence

Finally, we conducted mediation analyses to test whether NAcc R2’ may partially account for age-related changes in connectivity in the observed frontostriatal connections. Given the significant interaction effects between context and R2’ in two of the three ROI pairs, we conducted mediation analyses separately for reward- and resting-state contexts. We found that NAcc R2’ significantly mediated the relationship between age^-1^ and ventral ACC – NAcc connectivity in both reward-(Fig. 4C; *β* = 2.58, 95% CI [1.42, 3.95], *p* < .001) and resting-state (Fig. S2B; average indirect pathway: *β* = 1.36, 95% CI [.58, 2.26], p<.001). Similarly, NAcc R2’ significantly mediated age-related decreases in sACC – NAcc connectivity across both contexts (reward: Fig. 4C; *β* = 3.18, 95% CI [1.84, 4.74], *p* = .<.001; resting-state: Fig. S1B; β = 1.51, 95% CI [0.69, 2.47], p < .001). For the posterior medial OFC – NAcc pair, NAcc R2’ significantly mediated the relationship between age and connectivity in the reward-state (Fig. 4C; *β* = 1.48, 95% CI [.60, 2.57], *p* < .001), however mediation analyses were not conducted for resting-state, given the lack of association between connectivity and R2’ (see previous section). For simplification, values in Fig. 3C reflect only reward-state connectivity, and mediation analyses for resting-state connectivity are shown in Fig. S1B.

## Discussion

In this study, we used longitudinal, multimodal neuroimaging data to investigate the role of striatal tissue iron, a correlate of striatal DA neurophysiology, in the maturation of frontostriatal pathways through adolescence. We examined age-related changes in functional connectivity between the vmPFC and the NAcc at rest and during a reward state, and determined whether R2’ mediated developmental changes in connectivity. We found age-related decreases from adolescence to adulthood in the strength of functional connectivity between the NAcc and distinct nodes within the vmPFC that support reward processing (Fig. 4A), namely the vACC, subgenual cingulate, and the posterior medial OFC. Further, we show that NAcc tissue iron concentration affects individual variability in the strength of connectivity among this network (Fig. 4B), and significantly mediated developmental changes in vmPFC – NAcc connectivity (Fig. 4C). To our knowledge, this is the first study demonstrating that striatal tissue properties linked to DA function play a key role in the specialization of frontostriatal reward pathways throughout adolescence.

### Decreased vmPFC – NAcc Connectivity May Reflect a Developmental Shift in Reward Sensitivity Through Adolescence

Our results showing significant developmental decreases in vmPFC – NAcc coupling, evident in both reward and rest contexts, are in agreement with previous resting state and task-based fMRI studies showing developmental decreases in PFC – NAcc connectivity (Barnes et al., 2010; Fareri et al., 2015; Porter et al., 2015; Supekar et al., 2009; Van Duijvenvoorde et al., 2016; Van Duijvenvoorde et al., 2019, but see Van Den Bos et al., 2012 who showed increases in task-related connectivity between late childhood and early adolescence). These changes may be influenced by changes in DAergic processing and may contribute to known changes in adolescent reward processing. Prominent neurodevelopmental models propose that heightened reward-driven behavior, sensation seeking, and risk-taking during adolescence arises due to a relative predominance of affective systems (e.g., striatal DA) over cognitive control systems (e.g., PFC) during adolescence (Shulman et al., 2016). Specifically, the driven dual-systems model (Luna & Wright, 2016; Shulman et al., 2016) proposes that the adolescent peak in dopaminergic signaling within the NAcc predominates over new access to prefrontal executive systems, supporting sensation seeking in adolescence, which often reflects high level planning, but in service of obtaining immediate rewards (Luna & Wright, 2016). Our findings concur with earlier studies of structural corticostriatal connectivity supporting heightened influence of limbic corticostriatal circuitry in adolescence that wanes into adulthood. Evidence of developmental changes in the relative neural contributions of cognitive and affective systems are reflected in Larsen et al., 2017, who showed that in convergence zones in the striatum, the structural integrity of white matter projections originating from cortical limbic (affective) systems decreased with age, while those from cognitive control systems remained stable, resulting in a shift to a predominance of cognitive control inputs into adulthood, corresponding to decreases in the effect of rewards on behavior into adulthood (Larsen et al., 2017). Thus, the developmental decrease in connectivity observed in the current study (Fig. 4A) may be supported by decreased integrity of structural pathways linking limbic cortex to the striatum. Together, these findings may represent a mechanism that facilitates developmental decreases in exploratory drive and reward seeking behavior into adulthood (Larsen & Luna, 2018; Monica Luciana et al., 2012; Spear, 2000; Stansfield & Kirstein, 2006).

### Understanding Development of Reward Pathways through Adolescence

Central to the brain’s reward circuit are the NAcc and midbrain dopaminergic systems including the VTA and substantia nigra (Kelley & Berridge, 2002; Schultz et al., 2000; Wise, 2004). Importantly, these regions do not receive direct sensory input (Haber & Behrens, 2014), thus information about environmental cues, action-value representations, and cognitive control demands comes in from diverse cortical inputs, including the vmPFC (Haber & Behrens, 2014). The vmPFC, particularly the OFC and the ACC, is central to decision-making, with the OFC sharing connections with sensory systems, limbic systems, and cognitive control areas, and the ACC sharing connections with emotional, attentional, and regions involved in feedback processing (Haber & Behrens, 2014). These two regions provide the main cortical afferents to the NAcc, where they converge with inputs from the amygdala, hippocampus, and midbrain dopaminergic nuclei (Jarbo & Verstynen, 2015). The main midbrain outputs are to the striatum and cortex, including the NAcc, OFC, and ACC. Importantly, there are no known predominant direct projections from the NAcc to the cortex (but see Hoops & Flores, 2017); instead, the NAcc projects back to the ventral pallidum and the VTA/SN (Heimer et al., 1982; Heimer & van Hoesen, 1979), and to prefrontal cortical areas via the thalamus as part of an integrated corticobasal ganglia – thalamocortical loop (Alexander et al., 1986). The NAcc also relays information to prefrontal cortical areas via the nucleus basalis in the basal forebrain, which provides the main source of cholinergic fibers to the vmPFC (Haber & Knutson, 2010; Záborszky et al., 1993; Záborszky et al., 1991). This reward/incentive circuit is embedded within the corticobasal ganglia system and integrates information across diverse cortical and subcortical regions to bias action selection and mediate goal-directed behaviors (Haber, 2016; Haber & Knutson, 2010). Enhanced connectivity between the vmPFC – NAcc (Fig. 4A) in early adolescence, in combination with adolescent peaks in DA availability, could reflect several possibilities, including; 1. Increased input from the vmPFC to the NAcc in early adolescence that decreases into adulthood (Brenhouse et al., 2008); or 2. Increased NAcc signaling to vmPFC via corticobasal ganglia – thalamocortical loops in adolescence that decreases into adulthood. Increased activity among either of these pathways could result in increased correlated activity between the vmPFC and NAcc in adolescence (Fig. 4A), and, given the role of these pathways in reward processing, could partially underlie adolescent peaks in reward sensitivity that decrease into adulthood.

### Understanding Developmental Changes in Striatal Neurobiology through Tissue Iron

Tissue iron has been linked to several aspects of cognitive function, predominantly in early development and aging, and more recently, to the development of cognitive function through adolescence (Larsen, et al., 2020). Iron deficiency in early development corresponds to reductions in catecholamine synthesis (Beard, 2008; Torres-Vega et al., 2012), abnormal striatal DA function, including decreased D1 and D2 receptor concentration and DAT functioning (Erikson et al., 2000, 2001; Jellen et al., 2013; Lozoff, 2011), and cognitive impairment (see Grantham-McGregor & Ani, 2001; McCann & Ames, 2007 for review). In late life, the reverse pattern emerges whereby abnormally elevated tissue iron concentrations are associated with neuropathological processes underlying various neurodegenerative diseases (Bartzokis et al., 2004; Ward et al., 2014; Zucca et al., 2017), in addition to general cognitive decline (Daugherty & Raz, 2015; Ghadery et al., 2015; Penke et al., 2012; Pujol et al., 1992). Though several lines of evidence converge on a role for abnormal iron content in cognitive and DA dysfunction in early- and late-life, less is known about how variability in tissue iron levels affect the development of cognitive function and specialization of underlying neural circuitry during adolescence. However, a recent study highlighted a supporting role for putamen tissue iron in the development of cognitive function throughout adolescence, whereby individuals with low tissue iron concentrations (relative to those with high tissue iron) showed poorer cognitive ability during the transition from adolescence to adulthood (Larsen, et al., 2020). These results suggest that lower levels of striatal tissue iron potentially impact critical neurodevelopmental processes through the adolescent period. Our findings follow a similar neurodevelopmental trend, in that variability in NAcc tissue iron affects the development of frontostriatal circuits underlying reward processing (Fig. 4B). Specifically, *lower* NAcc tissue iron (relative to one’s age cohort) is associated with *elevated* frontostriatal connectivity (Fig. 4B), while *higher* tissue iron is associated with *decreased* frontostriatal connectivity (Fig. 4B). Given that connectivity *decreased* through adolescence into adulthood, our findings could reflect protracted maturation of frontostriatal connectivity in individuals with relatively low tissue iron for their age cohort. In further support of this notion, we showed that NAcc R2’ significantly mediated age-related decreases in connectivity (Fig. 4C), suggesting that ongoing maturation of NAcc tissue iron through adolescence may be an important contributor to the development of frontostriatal connectivity. Finally, voxelwise analysis confirmed the negative association between R2’ and frontostriatal connectivity (Fig. S3B), and revealed that this association was largely restricted to the vmPFC, illustrating a specified role for NAcc tissue iron in the specialization of mesolimbic frontostriatal circuitry.

### Tissue Iron, Dopamine, and Plasticity in Adolescence

Tissue iron supports myelination (Todorich et al., 2009) and DA-related functions, representing two key neuroplastic mechanisms that could potentiate the developmental changes in connectivity observed in the current study, and network specialization throughout adolescence more generally (Benoit-Marand & O’Donnell, 2008; Larsen & Luna, 2018; Porter et al., 1999; Steullet et al., 2014; Tseng & O’Donnell, 2007). Developmental models have posited that dynamic changes occurring within the mesocortical DA system through adolescence may facilitate neuroplasticity and specialization within cortico-striatal circuits, supporting maturation of cognitive function into adulthood (Kalsbeek et al., 1988, 1989; Larsen & Luna, 2018; Porter et al., 1999). Specifically, studies have shown pubertal peaks in D1 and D2 receptor density (Andersen et al., 1997; Tarazi et al., 1998a; Teicher et al., 1995, but see Matthews et al., 2013 & Sturman & Moghaddam, 2012, who did not find differences in adolescents relative to adults), DA concentration (Giorgi et al., 1987; Rao et al., 1991), and DAT levels (Moll et al., 2000; Tarazi et al., 1998b). Furthermore, adolescent increases in DA concentrations and the density of DA inputs to the PFC (Benes et al., 2000), axonal sprouting of DA axons from the NAcc to the PFC (Hoops & Flores, 2017), and the number of projections from the PFC to the NAcc (Brenhouse et al., 2008; Larsen & Luna, 2018) provide further mechanistic basis for heightened striatal functioning in adolescence. Although *in vivo* evidence of developmental changes in DA function in humans has been limited, we recently showed that while presynaptic vesicular DA, measured with PET [^11^C]dihydrotetrabenazine (DTBZ; (Brenhouse et al., n.d.) is developmentally stable by 18 years of age, D2/D3 availability, measured with PET [^11^C] Raclopride (RAC), continues to decrease in young adults (18-30 years of age; Larsen et al., 2020), potentially reflecting continued pruning of receptors into early adulthood. Enhanced DA function in adolescence and its effects on frontostriatal circuitry may facilitate reward- and sensation-seeking behavior in adolescence, which is adaptive for experience-dependent plasticity and specializing neurobiological pathways required for adult like levels of cognitive function (Larsen & Luna, 2018; Luciana et al., 2012; Spear, 2000). Thus, developmental decreases in DAergic function and innervation could be associated with dampening of vmPFC – NAcc coupling, as the circuit becomes refined through specialization.

## Conclusions

This study provides novel *in vivo* human evidence that DA-related striatal neurophysiology contributes to the specialization of frontostriatal circuitry through adolescence, potentially underlying known decreases in reward-driven sensation seeking into adulthood. Understanding neural mechanisms underlying normative adolescent specialization of critical circuitry can inform the emergence of psychopathologies such as substance abuse (Dionelis et al., 2019; Ernst & Luciana, 2015; Luciana et al., 2018), mood disorders (Davey et al., 2008; Diehl & Gershon, 1992), and schizophrenia (Howes et al., 2012; Paus et al., 2008), which emerge during this period and have been associated with impairments in frontostriatal systems. Here, we take a fundamental first step to characterizing a role for DA-related striatal neurobiology in the developmental specialization of frontostriatal circuitry in a large, longitudinal sample of healthy adolescents.

## Supporting information

Supplemental Methods & Results

## Acknowledgements

We thank the University of Pittsburgh Clinical and Translational Science Institute (CTSI) for help in recruiting participants. This work was supported by Grant Number 5RO1MH080243-07 from the National library of Medicine, National Institutes of Health, and the Staunton Farm Foundation.

