## Supplemental Methods & Results for "Dopamine-related striatal neurophysiology is associated with specialization of frontostriatal reward circuitry through adolescence"

*Functional Connectivity Analysis II: Whole-Brain Voxelwise Seeded Connectivity Analyses.*

*Voxelwise Functional Connectivity Analyses.* In order to examine the robustness of our a priori ROIs and provide unbiased estimates of all potential relevant frontostriatal connections, we repeated our connectivity analyses in a voxelwise framework. Bilateral NAcc ROIs were defined in the same manner as above, and represented the seed region for the whole-brain voxelwise analyses.

To assess functional connectivity across contexts, we extracted the timeseries from bilateral NAcc ROIs (as defined above, 3dDeconvolve in AFNI) and calculated voxel-level spatial maps from their correlations with voxels in a mask of all grey matter voxels in the frontal cortex (See Fig. S2) that was created from the MNI-152-09c template in AFNI (3dcalc in AFNI). The mask was created to include voxels anterior to $y = 31$ with a 50% or greater probability of being grey matter in the MNI-152-09c template. We restricted our analyses to a frontal mask as our hypotheses were centered around understanding developmental changes in frontostriatal circuitry. Voxelwise linear mixed-effects analyses were implemented in AFNI (3dLME; Chen et al., 2013) and testing was limited to voxels had full EPI coverage on all participants across all runs. To assess differences in connectivity as a function of age and context, we used a linear mixed-effects model as implemented in AFNI's 3dLME with state (rest, reward), age, sex, head motion, seed ROI hemisphere, laterality, and visit number as fixed-effects, and random intercepts where estimated for each participant. Multiple comparison correction for this analysis was performed using the intersection of voxelwise FDR correction ($q$ <0.05) and cluster size. AFNI's 3dClustsim program (with acf option) was used to determine cluster size threshold through a Monte Carlo simulation with parameters derived from mean spatial autocorrelation parameters from the residual time series output in 3dDeconvolve.

***Supplemental Results***

*NAcc – vmPFC Connectivity Decreases Throughout Adolescence*

Main effects of age were also observed in the vmPFC at the voxelwise level (Fig. S3A, Table S1), with consistent patterns of decreased NAcc – vmPFC connectivity with increasing age across both resting-state and reward-state contexts (Fig. S3A, Table S1). Fig. S4A shows clusters for a lenient voxel-wise threshold of $p<0.05$ (uncorrected) and $n > 36$ contiguous voxels, as full statistical voxelwise criteria was not met. Increases in connectivity with age were also observed at the voxelwise level, particularly in areas of the lateral PFC (Fig. S3A, Table S1).

*Decreased NAcc – vmPFC Connectivity is Associated with Higher NAcc Tissue-Iron*

In confirmation of the ROI results, main effects of NAcc R2' were also observed at the voxelwise level (Fig. S4B & Table S1) in a large cluster largely restricted to the vmPFC (Fig. S3B & Table S1), whereby greater NAcc R2' values were associated with decreased connectivity. Fig. S3B shows clusters for a voxel-wise threshold of $p < .001$, FDR corrected ($q$ <.05), and $n > 36$ contiguous voxels.
