## Supplemental Figure 1 for "Dopamine-related striatal neurophysiology is associated with specialization of frontostriatal reward circuitry through adolescence"

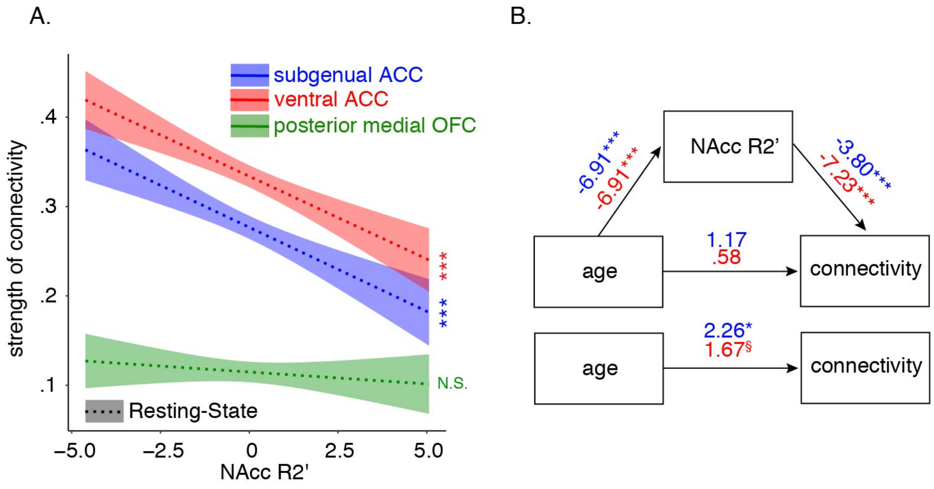


**Supplemental Figure 1.** Associations between nucleus accumbens R2' and resting-state connectivity. (B) Higher R2' (age-residualized) is associated with decreased connectivity between the nucleus accumbens and the ventral anterior cingulate and subgenual cingulate, but not the orbitofrontal cortex. In (B), mediation results for the nucleus accumbens – orbitofrontal cortex not shown due to lack of significant association between R2' and connectivity. In (A), the x-axis represents age-residualized estimates of R2', and shaded region represents 95% confidence interval. * $p_{\text{Bonferroni}} < .05$, ** $p_{\text{Bonferroni}} < .01$, *** $p_{\text{Bonferroni}} < .001$. ACC, anterior cingulate cortex, OFC, orbitofrontal cortex.
