## Supplemental Figure 2 for "Dopamine-related striatal neurophysiology is associated with specialization of frontostriatal reward circuitry through adolescence"

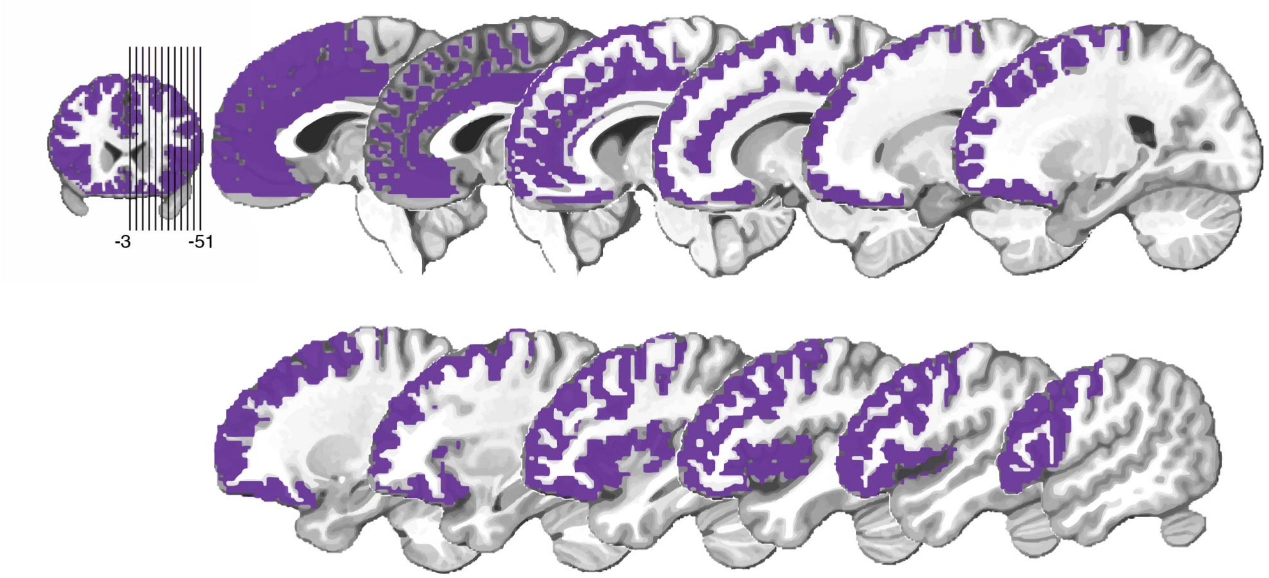


**Supplemental Figure 2.** Frontal mask used for voxelwise analyses. Voxelwise analyses were restricted to a frontal mask that was created using the MNI – 152 – 09c template in AFNI (3dCalc). Voxels anterior to y = 31 with a 50% or greater probability of being grey matter in the MNI – 152 – 09c template were included in the mask. The right hemisphere is depicted for visualization (x = -3 ranging to x = -51), and the inset reflects the axial slice location.
