## Supplemental Figure 3 for "Dopamine-related striatal neurophysiology is associated with specialization of frontostriatal reward circuitry through adolescence"

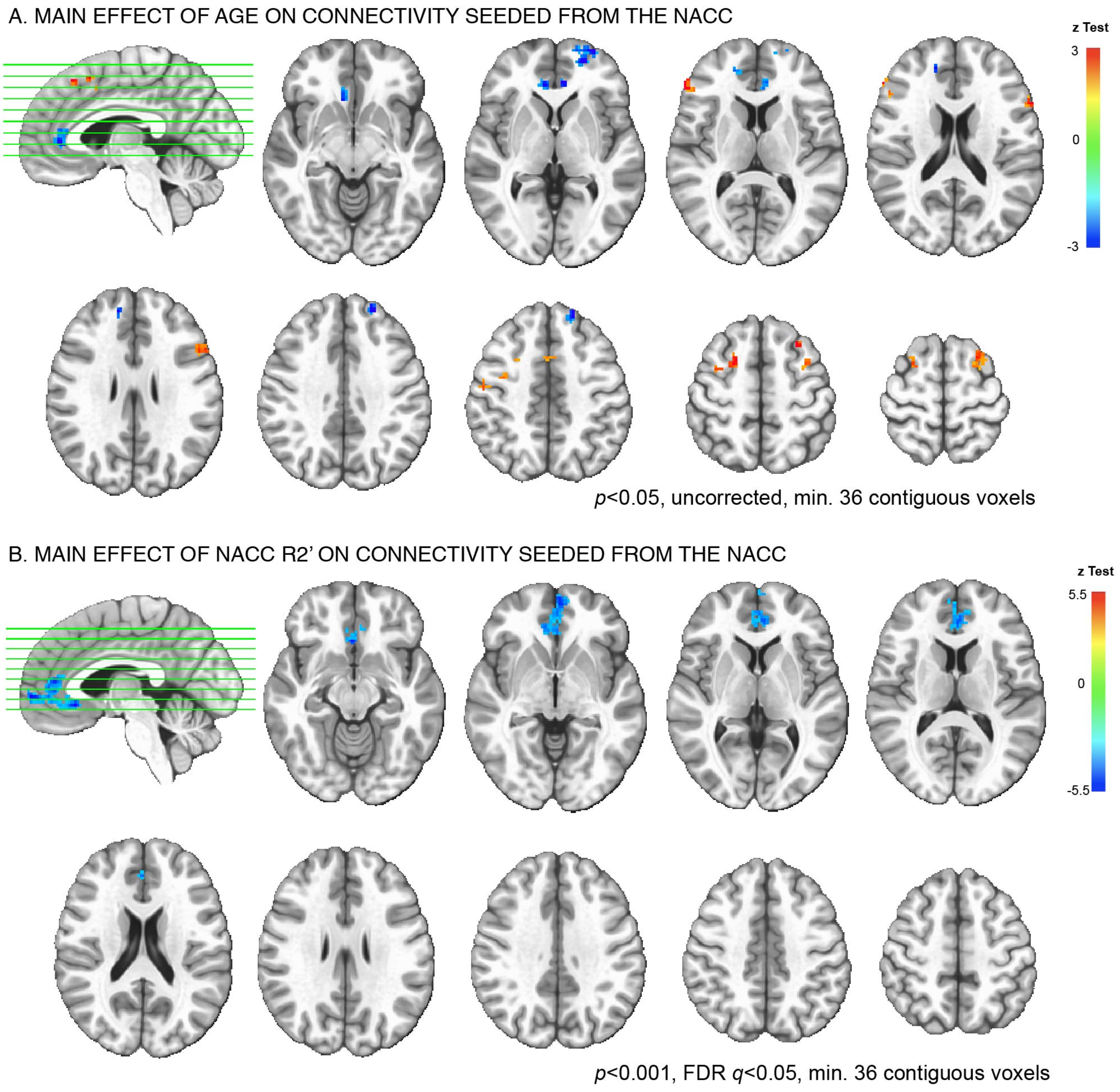


**Supplemental Figure 3.** Exploratory voxelwise analysis of nucleus accumbens – frontal cortex connectivity. (A) Main effects of age. Clusters of voxels (within the frontal mask in Fig. S3) showing significant age-related change (inverse age, $age^{-1}$) in functional connectivity with the nucleus accumbens. Clusters are shown for a lenient voxel-wise threshold of $p < .05$ and n > 36 contiguous voxels. (B) Main effects of nucleus accumbens R2' (age-residualized). Clusters of voxels showing significant change as a function of R2' in functional connectivity with the nucleus accumbens. Clusters are shown for a voxelwise threshold of $p < .001$, FDR corrected ($q < .05$), and n > 36 contiguous voxels. Color indicates effect size of the $age^{-1}$ (A) and R2' (B) terms, and sign indicates an overall increase (red) or decrease (blue) with increasing age and R2', respectively. NAcc, nucleus accumbens.
