## Supplemental Table 1 for "Dopamine-related striatal neurophysiology is associated with specialization of frontostriatal reward circuitry through adolescence"

| Test | Region | BA | Size of cluster | MNI Coordinates | | |
|---|---|---|---|---|---|---|
| | | | | Peak X | Peak Y | Peak Z |
| **Main effect of age** | R vACC | 24 | 218 | 1 | 43 | 5 |
| | L vmPFC | 14 | 39 | -10 | 43 | -8 |
| | R dACC | 32 | 41 | 15 | 25 | 28 |
| | L frontopolar | 10 | 104 | -24 | 57 | -6 |
| | R DLPFC | 6 | 417 | 26 | 6 | 51 |
| | L DLPFC | 6 | 301 | -31 | 15 | 63 |
| | R IFG | 44 | 76 | 59 | 13 | 17 |
| | L IFG | 44-45 | 111 | -61 | 18 | 17 |
| | R MFG | 46 | 57 | 56 | 36 | 10 |
| | L MFG | 46 | 99 | -43 | 36 | 19 |
| | R Insula | | 54 | 33 | 15 | -11 |
| | SMA | 6 | 48 | -3 | 15 | 54 |
| **Main effect of R2’** | R vmPFC | | 593 | 6 | 36 | -6 |

**Supplemental Table 1.** Clusters of voxels showing significant (A) age-related differences, (B) changes with R2’ in functional connectivity with the NAcc (visualized in Fig. S3). Note: BA, Brodmann area; MNI, MNI coordinates at peak, DLPFC, dorsolateral prefrontal cortex, vACC, ventral anterior cingulate cortex, IFG, inferior frontal gyrus, MFG, middle frontal gyrus, SMA, supplementary motor area, dACC, dorsal anterior cingulate, vmPFC, ventromedial prefrontal cortex, R, right, L, left.
